# Genomic and Epigenomic Relationships Capture Different Patterns of Genotype-by-Environment Interaction in Grapevine Clones

**DOI:** 10.64898/2026.11.05.764487

**Authors:** Hannah Robinson, Maximilian Schmidt, Paolo Callipo, Carlos A. Robles-Zazueta, Stephanie Brunner, Jessica Hintzsche, Ulduz Vafadarshamasbi, Johanna Frotscher, Timo Strack, Peter Crisp, Hans-Peter Piepho, Kai P. Voss-Fels

**Affiliations:** Department of Plant Breeding, Hochschule Geisenheim University, Geisenheim, Germany; Queensland Alliance for Agriculture and Food Innovation, Centre for Crop Science, The University of Queensland, St Lucia, Australia; School of Agriculture and Food Sustainability, Faculty of Science, The University of Queensland, St Lucia, Australia; Biostatistics Unit, Institute of Crop Science, University of Hohenheim, Stuttgart, Germany

## Abstract

Genotype-by-environment interaction (GEI) challenges selection in clonally propagated perennial crops by reducing the consistency of clone performance across years. In perennial crops, both genetic and epigenetic variation are likely to contribute to complex trait variation among clones. However, the extent to which each molecular source contribute to GEI remains poorly understood. Here, we used UMR-seq, an assay that recovers SNPs and unmethylated- region (UMR; DNA methylation) profiles from a single library, to generate paired genomic and epigenomic profiles for 206 Riesling (*Vitis vinifera* L.) clones phenotyped for yield and juice quality traits across nine growing seasons. We then partitioned the contribution of each omic layer to clonal variation and GEI, and examined the genomic distribution of their effects. In models without explicit GEI, environmental and residual effects were the largest source of phenotypic variance and the clonal main effect was modest. Genomic relationships accounted for most of the clonal variance in four of five traits, and adding an epigenomic relationship matrix (ERM) to the genomic relationship matrix (GRM) had little effect. In contrast, when GEI effects were modelled using factor-analytic covariance structures, the epigenomic layer captured a substantial proportion of GEI variance and exceeded the genomic contribution in four of five traits. The two omic layers differed not only in the amount of GEI captured, but also in the pattern of clone re-ranking across years. When back-solved effects were aggregated into genomic windows, the largest genomic and epigenomic effects tended to fall in different regions, with overlap no greater than expected by chance. Together, these results show that genomic and epigenomic relationships capture complementary components of year-to-year variation in clone performance, with the epigenomic contribution expressed primarily through GEI. These findings provide a basis for investigating the molecular drivers of GEI and for testing whether genomic and epigenomic information can inform selection for stable clonal performance across variable environments.

## Introduction

Phenotypic expression of complex traits reflects genetic potential, environmental conditions, management practices, and the interactions among these components (Falconer & Mackay, 1996). This genotype-by-environment-by-management interaction remains one of the most persistent challenges in plant breeding, limiting the reliable transfer of selection decisions across years and production environments. For simplicity we focus herein on genotype-by- environment interactions (GEI) that generally occur in two forms. First scale-type interaction, in which genotype rankings are conserved across environments but differ in size, and are largely less problematic in breeding. In contrast crossover interactions, where genotypes re- rank and in some cases reverse ranking, challenges selection across environments and complicate the identification of broadly adapted genotypes. As climate change intensifies, weather extremes are expected to become more frequent and intense (Seneviratne et al., 2021). Models that quantify and interpret GEI across current and future production environments are thus increasingly important for improving the reliability of selection decisions and sustaining crop productivity (Cooper et al., 2021; Cooper & Messina, 2023).

Grapevine (*Vitis vinifera* L.) is a particularly relevant system in which to examine this challenge. Grape production in many established regions is already approaching its climatic limits, with more frequent heat, drought and extreme weather events reducing yields and compromising fruit quality (van Leeuwen et al., 2024). Yield and juice quality traits are highly sensitive to interannual environmental variation, making GEI a persistent constraint on genetic improvement in perennial grapevine. In addition, grapevine is highly heterozygous and cannot be reproduced true-to-type from seed, so established varieties have been maintained through vegetative propagation for centuries to preserve their genetic integrity. Together with a long juvenile phase, this extends breeding cycles to as much as 25 years and severely limits the rate of genetic gain (van Leeuwen et al., 2024; Schmidt et al., 2025). Predictive modelling can help accelerate this gain, but its value depends on accurately predicting GEIs. Yet, existing studies in grapevine have largely been conducted in single environments or with simplified GEI terms that do not explicitly model the covariance structure of genotype performance across years and locations (Brault et al., 2021; Brault et al., 2022a; Brault et al., 2022b; Flutre et al., 2022; Brault et al., 2024).

For established grapevine varieties, genetic improvement can be achieved through clonal selection, in which clones carrying naturally occurring or induced variation are selected for superior trait performance. This allows gains to be made while preserving varietal identity. Over centuries of vegetative propagation, clones accumulate intra-varietal diversity through somatic mutations and epigenetic changes, creating mitotically heritable phenotypic differences in important traits (Carbonell-Bejerano et al., 2017; Vondras et al., 2019; Xiao et al., 2025; Callipo et al., 2026). Epigenetic variation describes differences in gene regulation that arise without modifying the DNA sequence (Berger et al., 2009). The principal forms are cytosine methylation, histone modifications and non-coding RNA-mediated silencing, which together shape chromatin structure and the accessibility of genetic information. In plants, cytosine methylation is the most commonly profiled form of epigenetic variation, where it regulates gene expression and contributes to genome stability through transposable-element silencing (Zhang et al., 2018).

The contribution of epigenetic mechanisms to clonal differentiation in grapevine, and in clonally propagated crops more broadly, is only beginning to be characterised. Plant DNA methylation occurs in three sequence contexts with distinct inheritance dynamics. CG and CHG methylation are comparatively stable and maintained through DNA replication, whereas CHH methylation is re-established de novo and is generally more dynamic. Vegetative propagation bypasses meiosis, allowing methylation states to be transmitted mitotically across successive propagation cycles and growing seasons (Douhovnikoff & Dodd, 2015; Zhang et al., 2018). Methylation differences can also arise through spontaneous epimutation and persist independently of changes in DNA sequence or environmental cues. Consistent with this potential for mitotic inheritance, an early study demonstrated that methylation-based markers could discriminate among lineages of Pinot Noir clones (Ocaña et al., 2013). More recent genome-wide profiling showed that CG and CHG methylation polymorphisms reconstructed clonal relationships that closely mirrored somatic SNP-based relatedness (Callipo et al., 2026). These findings indicate that a component of epigenetic variation arising during clonal propagation is at least partly maintained across vegetative generations.

Alongside this stable component, DNA methylation is also environmentally responsive. Unlike somatic mutations, which are stably propagated once established, methylation states can be established or removed rapidly in response to external stimuli (Crisp et al., 2016; Callipo et al., 2025). In Shiraz, methylation profiles differentiated vines grown in different vineyards despite limited genetic differentiation (Xie et al., 2017), while in Malbec the extent of methylation differentiation between contrasting vineyard environments varied among clones, indicating that environmental responses can themselves be clone dependent (Varela et al., 2021). When cuttings of a single Malbec clone from two contrasting vineyards were grown in a common vineyard, differences in global methylation levels were lost within two years, whereas vineyard-of-origin differences in methylation patterns diminished more gradually (Varela et al., 2024). Evidence linking such methylation variation directly to measured trait differences nevertheless remains limited, with Varela et al. (2024) detecting no association between methylation and phenotype in the same vines. The coexistence of mitotically stable and environmentally responsive methylation variation therefore provides a strong rationale for testing whether methylation data capture trait variation and environmental response beyond those captured by genomic data.

Testing these questions requires genomic and epigenomic profiles for populations large enough for quantitative genetic analysis. The use of epigenomic data in predictive breeding has remained limited, largely because such data are costly to generate. In grapevine, methylation has mostly been profiled by bisulfite or enzymatic conversion sequencing, or by reduced- representation methods that sample only a small fraction of the genome (Schmidt et al., 2025). UMR-seq takes a different approach. In plants, unmethylated regions (UMRs) are largely stable across vegetative tissues and mark expressed genes and their cis-regulatory space (Crisp et al., 2020). Building on this, UMR-seq uses a combination of methylation-dependent restriction enzymes to fragment unmethylated DNA. It provides a genome-wide readout of UMRs in both genic and intergenic regions, together with SNPs called from the same reads (Putland et al., 2026). A single low-input library therefore generates paired genomic and epigenomic profiles.

Whether genomic and epigenomic variation capture distinct or overlapping patterns of GEI remains unclear, particularly in grapevine, and requires joint modelling of both layers. Consistent with this, combining epigenomic and transcriptomic data with genomic markers in perennial ryegrass explained more phenotypic variance than any single-omic model (Malinowska et al., 2022). Across crops more broadly, such gains appear context-dependent, for example in a benchmarking study across rice and maize, multi-omic integration increased the variance captured in all three datasets but improved prediction accuracy in only one (Montesinos-López et al., 2025). Here, we applied UMR-seq to 206 clones from the Riesling collection at Hochschule Geisenheim University (HGU), in which clonal differences in yield and juice quality traits are repeatable across seasons (Robinson et al., 2025a). To our knowledge, this is among the largest paired genomic and epigenomic datasets for clones of a single grapevine variety. We hypothesise that SNP and methylation data capture partially independent sources of trait variation among these clones, and that the two omic layers differ in their contributions to clonal variation and GEI. We test this through three objectives: (i) characterise genomic and epigenomic differentiation among clones and quantify each omic layer’s contribution to clonal variance; (ii) quantify their contributions to GEI and determine whether they capture distinct patterns of environmental response; and (iii) explore the genomic distribution of genomic and epigenomic effects and determine whether effect magnitude is associated with deviation from the seasonal response.

## Materials and Methods

### Plant material and experimental design

The study comprised 206 Riesling (*Vitis vinifera* L.) clones selected from the Riesling collection established at Hochschule Geisenheim University, Geisenheim, Germany, and described in full in Robinson et al. (2025a). Clones were selected to provide broad representation of the diversity present within the larger collection. The wider Riesling collection was assembled from historic vineyards across Germany, France, Austria, and Luxembourg, with individual clones originally selected for advantageous agronomic characteristics. The selection location of each clone is listed in Supplementary Table S1. Following virus screening and varietal confirmation, the clones were propagated from single- bud cuttings, grafted onto phylloxera-resistant rootstocks, and established under standardized management in the experimental vineyard. Rootstock assignment and further details of clone provenance, propagation, and viticultural management are reported in Robinson et al. (2025a).

### Phenotypic trait observations

Riesling clones were evaluated for yield and juice quality traits across harvest seasons from 2009 to 2023. Five seasons with fewer than 30 scored plots (2009, 2010, 2013, 2014 and 2023) were excluded because season-specific variance components could not be estimated reliably, resulting in 206 clones measured across nine seasons (2012 and 2015 to 2022). Five traits were examined: total berry yield (g m^−2^), total soluble solids (°Brix), total acidity (g L^−1^), tartaric acid (g L^−1^) and malic acid (g L^−1^). Phenotypic measurement protocols followed Robinson et al. (2025a). Briefly, must quality traits were determined from pressed juice by Fourier- transform infrared spectroscopy using a Winescan FT2 spectrometer (FOSS, Hillerød, Denmark) with an in-house grape must calibration. Outlier removal and raw data curation procedures also followed Robinson et al. (2025a). To explore phenotypic variation prior to omic modelling, best linear unbiased estimators (BLUEs) of clone means were obtained for each trait. BLUEs were derived from a linear mixed model with clone and year fitted as fixed effects, field and year × field as random effects, and year-specific residual variances. Clonal repeatability was estimated for each trait by fitting clone as an independent random term, rather than fixed, in the same linear mixed model to estimate best linear unbiased predictors (BLUPs). Clonal repeatability was calculated following the generalised heritability approach of Cullis et al. (2006) as:

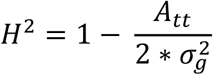

Where *A_tt_* is the mean variance of a pairwise difference between clone BLUPs and σ^2^_*g*_ is the clonal variance.

### Plant tissue sampling

Plant tissue was collected from field-grown vines between 14 and 20 May 2021, at the phenological stage BBCH 15 (five leaves unfolded; Lorenz et al., 1995). For each clone, three newly fully expanded leaves were sampled from the middle section of the cane. Each leaf was collected from a different vine, resulting in one leaf sampled from each of three replicate vines per clone. Immediately after collection, samples were flash frozen in liquid nitrogen and transported to the laboratory, and stored at −70 °C until DNA extraction.

### DNA extraction and UMR-seq library preparation and sequencing

Pooled leaves were ground to a fine powder in liquid nitrogen. DNA was extracted from 150 mg of homogenised leaf material using the innuPREP Plant DNA Kit (Analytik Jena, Jena, Germany) following the manufacturer’s Protocol 2 (Extraction using Lysis Solution OPT). Lysis was performed at 65 °C for 60 min. DNA was eluted in demineralised water.

UMR-seq libraries were prepared following the method developed by Putland et al. (2026), with the modifications described below. For each Riesling clone, 250 ng of genomic DNA was digested with the methylation-dependent restriction enzymes MspJI and FspEI to preferentially deplete methylated DNA and enrich for unmethylated genomic regions. Digestion was performed in a final reaction volume of 30 µl containing 1x CutSmart buffer, 2.5 U each of MspJI and FspEI, and 1x Enzyme Activator (New England Biolabs, Ipswich, MA, USA). Reactions were incubated at 37 °C for 2 h, followed by enzyme inactivation at 65 °C for 20 min. Digested DNA was purified using SPRIselect-based size selection (Beckman Coulter, Lane Cove, NSW, Australia) with the modifications described by Crisp et al. (2026) and summarized below. A 25 µl aliquot of each digest was purified using a 0.7x bead-to-digest ratio by adding 17.5 ul of beads, to remove fragments less than 200 bp, mixing steps were performed by slowly pipetting approximately 20 times, and DNA was eluted in 21 µl of ultrapure water with a 5 min incubation before transferring the eluate containing the unmethylated DNA fragments to a new storage tube. Purified DNA was quantified using a Qubit fluorometer with the 1x High Sensitivity assay (Life Technologies, Eugene, OR, USA). Digested and purified DNA samples were also assessed by 1% agarose gel electrophoresis to evaluate digestion and purification quality. Purified DNA was diluted to 0.2 ng µl-1, and 1 ng was used for library preparation.

Libraries were prepared using the Nextera XT DNA Library Prep system (Illumina, 2019), with the modifications for methylation-dependent restriction-digested DNA described by Crisp et al. (2026) and summarized below. For tagmentation, 1 ng of purified DNA was used, while the volumes of all other reaction components were reduced to half those specified in the standard Nextera XT protocol. Tagmentation was performed at 55 °C for 5 min, followed by neutralization at room temperature for 5 min. Tagmented DNA was subsequently amplified in a total reaction volume of 32.5 µl, comprising 15 µl of tagmented DNA, 5 µl of each adapter and 7.5 µl of Nextera PCR Master Mix. PCR amplification was performed using the standard Nextera XT thermocycling conditions (Illumina, 2019) for 12 cycles. Following amplification, libraries were purified using SPRIselect beads according to the Nextera XT protocol, with 45 µl of beads added to 27.5 µl of amplified library. Purified libraries were quantified using the Qubit 1x High Sensitivity assay. All indexed samples were then pooled such that each sample contributed an equal mass to the final library pool. Final library pools were sequenced on an Illumina NovaSeq S4 platform at the Australian Genome Research Facility (AGRF) using 150- bp paired-end sequencing.

### UMRseq read trimming and alignment

Raw paired-end reads from UMRseq libraries were trimmed with Trimmomatic v0.39 (Bolger et al., 2014) to remove Nextera adapters and low-quality bases. Surviving read pairs were aligned to the V. vinifera PN40024.v4 reference genome (Velt et al., 2023) with BWA-MEM2 v2 (Vasimuddin et al., 2019), coordinate-sorted with samtools (Danecek et al., 2021), and PCR duplicates removed with Picard MarkDuplicates (Broad Institute, 2019). Two downstream pipelines were then run on the deduplicated BAMs: SNP calling and coverage-based differential methylation. Full commands and parameters are available at Schmidt (2026).

### SNP-calling pipeline

Per-sample genotype likelihoods were computed with GATK HaplotypeCaller in GVCF mode, combined per chromosome with GenomicsDBImport, and jointly genotyped with GenotypeGVCFs (McKenna et al. 2010; Poplin et al. 2018). Raw variants were hard-filtered following GATK Best Practices (QD < 2.0, FS > 60.0, MQ < 40.0, MQRankSum < −12.5, ReadPosRankSum < −8.0; Poplin et al., 2018). Genotype-level filtering with VCFtools (Danecek et al., 2011) retained sites with per-genotype depth ≥ 3, present in ≥ 50% of samples, mean depth ≥ 5, and minor-allele frequency ≥ 0.05 across the 217 sequenced samples. Per- chromosome VCFs were concatenated into a whole-genome variant set.

### Coverage-based differential methylation pipeline

Since UMR-seq selectively enriches unmethylated genomic regions, the local read coverage of a sample reflects its methylation state at that locus, and differential coverage between samples can be used as a proxy for differential methylation. We therefore developed a parallel coverage pipeline based on the same deduplicated BAMs. For each sample, deduplicated alignments were filtered to retain uniquely mapped, properly paired reads, converted to fragment intervals with bedtools (Quinlan & Hall, 2010), and fragments longer than 1 kb or on organellar contigs were discarded. A common window set was defined from pooled coverage across all samples, excluding windows below 10x pooled coverage or shorter than 50 bp. Counts were further normalized to an FPKM-like metric, computed as (reads × 10⁹) / (total sample reads × window length), with windows yielding values ≤ 1 set to zero.

### Omic data curation and relationship matrices construction

Following matching to phenotypic records, 206 clones were retained, allele frequencies were recomputed within this subset, and markers were filtered again at MAF ≥ 0.05. To summarise how variable each methylation feature was among clones, we computed its Shannon entropy, *H* = −Σ*_b_ p_b_* log_2_ *p_b_*, where *p_b_* is the proportion of clones whose normalised coverage at that feature falls into bin *b* of 20 equal-width bins spanning that feature’s observed range across clones. Entropy approaches zero when nearly all clones fall into the same bin and reaches its maximum (log_2_ 20 ≈ 4.32) when clones are spread evenly across bins. Genome-wide additive relatedness among clones was modelled using a VanRaden (2008) genomic relationship matrix (GRM). SNP genotypes were coded as allele dosages and centred by marker allele frequencies, such that *Z_ij_* = *S_ij_* − 2*p_j_*, where *S_ij_* is the genotype score for clone *i* at marker *j* and *p_j_* is the corresponding allele frequency. The GRM was then calculated as

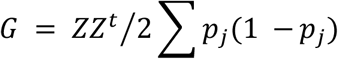

placing genomic relationships on an additive-genetic scale. An equivalent epigenomic relationship matrix (ERM) was constructed as

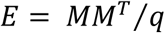

where *M* is the mean-centred *n* × *q* matrix of continuous methylation values and *q* is the number of methylation features. To address collinearity between the GRM and ERM, the methylation matrix was adjusted for the first 10 genomic principal components, where each locus was regressed on the first 10 principal components of the GRM, and the residuals were centred, scaled, and used to recompute the ERM using the formulation above. Since only the leading genomic axes were removed, this adjustment reduces but does not fully remove collinearity with the GRM; for brevity, we refer to the resulting matrix as the orthogonalised ERM. All relationship matrices were placed on a common scale using the average semi- variance (ASV) framework (Piepho, 2023; Feldmann et al., 2022). Each matrix *K* was transformed as *PKP*, where *P* = *I* − *n*^−1^11*^T^*, and scaled such that *tr*(*PKP*)⁄(*n* − 1) = 1. Variance components estimated under this scaling are expressed as half the average variance of a difference between clones and are therefore directly comparable across omic kernels. Agreement between relationship matrices was quantified as the Pearson correlation coefficient of corresponding off-diagonal elements. Associations between relationship matrices and clone passport data were tested by permutational multivariate analysis of variance (PERMANOVA; 999 permutations) on distances derived from each relationship matrix (d²ᵢⱼ = Kᵢᵢ + Kⱼⱼ − 2Kᵢⱼ), for region of selection (regions represented by fewer than five clones pooled), generation and vineyard field. Tests were applied to the GRM, the orthogonalised ERM and an ERM computed from the unadjusted methylation features.

### Multi-kernel Non-GEI mixed models

All models were fitted using ASReml-R v4.2 (Butler et al., 2023) via restricted maximum likelihood (REML). A baseline model incorporating both omic layers was fitted for each trait:

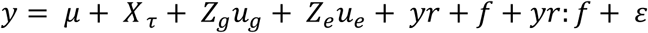

where *y* is the vector of phenotypic observations across all years and *μ* is the overall fixed intercept. *τ* contains the fixed effects of field and virus status. 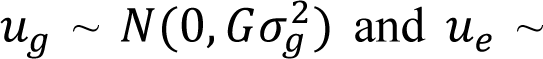 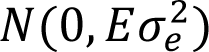 are vectors of additive genomic and epigenomic effects with corresponding relationship matrices *G* (GRM) and *E* (ERM). Additional random effects include *yr*, *f*, and *yr*: *f* which are *year*, *field*, and *year* × *field*, with *y*: *f* modelled as a diagonal variance structure. Residuals were modelled with year-specific variances, such that *ε* ∼*N*(0, ⊕*_t_ σ*_2_ *ε*, *t I_nt_*), modelling heterogeneous residual variances across (*t*) years. A sparse field design precluded inclusion of spatial row and column terms in the residual structure. Non- additive kernels were evaluated in preliminary analyses (Supplementary Methods). Across all non-additive kernels and traits, the largest gain in log-likelihood over the additive GRM + ERM model was 0.82, below the threshold for significance in a boundary likelihood-ratio test. Non- additive kernels were therefore not retained in any model reported herein.

Models were specified in parallel, differing only in the relationship matrices included, so that differences in captured clonal variance could be attributed to the relationship matrices rather than model structure. All models shared the same fixed effects and environmental terms, so that restricted log-likelihoods and AIC were directly comparable. Four models were fitted per trait: M1 (reference): 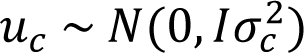, an independent clonal main effect with no relationship matrix; M2 (GRM): 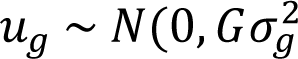, a genomic only effect; M3 (GRM + ERM): *u_g_* + *u_e_*, as in the baseline model;

M4 (GRM + ERM + clone): M3 plus *u_c_*, to test whether clonal variance escaped both kernels. For the kernel models, the clonal variance used to compute clonal repeatability was the sum of the kernel variance components 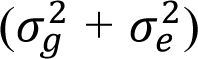, which under ASV scaling equals half the average variance of a difference between two clones.

### GEI mixed models and back-solving of locus effects

To characterise GEI, the baseline model was extended to allow omic-layer-specific clone effects to vary across years. The full model for each trait was:

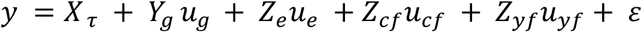

where *τ* contains the fixed effects of year, field and virus status; *u_g_* and *u_e_* are the vectors of genomic and epigenomic clone-by-year effects; 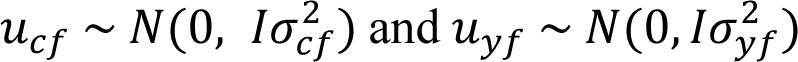 and 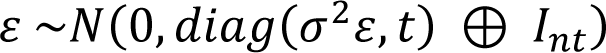 are the clone-by-field and year-by-field effects; and *ε* ∼*N*(0, *diag*(*σ*^2^*ε*, *t*) ⊕ *I_nt_*) has year- specific residual variances. For each trait, clone-by-year effects were modelled separately for each omic layer using a factor-analytic (FA) covariance structure across years (Smith et al., 2001), such that:

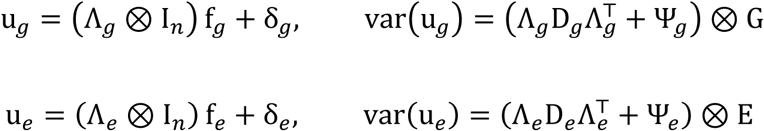

where Λ is a *t* × *k* matrix of year loadings with orthonormal columns **Λ**^T^**Λ** = I*_k_*; *f* is the vector of clone scores on the *k* factors, with score variances D = diag(*d*_1_, … , *d_k_*) ordered so that *d*_1_ > *d*_2_ > ⋯ > *d_k_*; and δ holds the year-specific deviations, with specific variances **Ψ** = diag(*ψ*_1_, … , *ψ_t_*). Subscripts *g* and *e* denote the genomic and epigenomic layers, and *G* and *E* the corresponding relationship matrices. The FA order and other trait-specific variations in the fitted terms are listed in Supplementary Table S2.

Correlations among years were derived separately for each omic layer from the fitted FA covariance matrices and used to characterise scale-type and crossover GEI for each layer. To examine where in the genome the effects captured by each layer arise, year- and omic-specific BLUPs were extracted from the GEI model for each clone, and locus-level effects were recovered by back-solving from these BLUPs using a dual-form ridge regression:

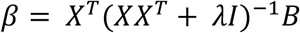

where *X* is the *n x p* matrix of SNP allele dosages (0, 1, 2) or orthogonalised methylation features, *B* is the *n x t* matrix of clone BLUPs across *t* years for the corresponding omic layer, *I* is the *n x n* identity matrix. Due to *XX^T^* being singular or ill-conditioned (for the orthogonalised methylation features, its rank is at most *n* − 11 after column-centring and the removal of ten genomic principal components; for the uncentred SNP dosages it is at most *n*, but the matrix is ill-conditioned when p ≫ n), a small ridge penalty (λ = 0.1, negligible relative to the diagonal elements of *XX^T^*) was applied for numerical stability. Back-solved effects are expressed per unit of each layer’s feature coding (allele dosage for SNPs; standardised coverage for methylation features) and are spread across a different number of features in each layer, so their absolute magnitudes are not comparable between layers. Comparisons between layers were therefore based on within-layer ranks or on values scaled to each layer’s 95th percentile. The resulting matrix is a *p x t* matrix of locus-level effects β per year.

### Window-level analysis of back-solved effects

Since individual back-solved locus effects can be small and noisy, effects were summarised at a regional scale, by aggregating into common 25 kb sliding windows advanced in 10 kb steps, so that consecutive windows overlapped by 15 kb. The 25 kb window balanced positional resolution against the number of features per window, and the 10 kb step avoided splitting effects arbitrarily at window boundaries. Windows were defined on the same coordinates for both layers and retained for a layer if they contained at least five features of that layer (SNPs for the genomic layer; methylation features for the epigenomic layer), because per-feature means over very few features are unstable.

Within each window and year, the window effect was calculated as the mean of the absolute back-solved effects of the features in that window. Absolute values ensure that effects of opposite sign within a window do not cancel. The mean was chosen rather than the sum, as back-solving assigns a non-zero effect to every feature, therefore summed scores largely reflected feature density. Preliminary analyses showed summed window scores were almost perfectly correlated with feature count and varied little between traits (Spearman ρ between yield and tartaric acid window rankings: 0.99 for SNPs and 0.92 for methylation features). The window effect therefore measures the average effect per feature in a region, and may understate regions in which many features each contribute modestly. Effect magnitude was defined as the mean window effect across the nine years.

Seasonal deviation was quantified following Eberhart and Russell (1966). For each window, the nine yearly window effects were regressed on an environmental index, defined as the mean window effect across all windows in that year, and seasonal deviation was calculated as the root mean square deviation (RMSD) of the yearly effects from the fitted line. Seasonal deviation therefore measures how far a window departs from its own linear response to the seasonal index, thus a window whose effect rises and falls in proportion to the index has low seasonal deviation even if its effect changes considerably between years. It corresponds to the deviation-from- regression component of Eberhart and Russell (1966), i.e. stability in the sense of predictability (Lin et al., 1986), and is distinct from crossover GEI, which was assessed from the between- year correlations of the factor-analytic models. Since window effects are based on absolute locus effects, seasonal deviation reflects changes in the size of a window’s effect across years and does not capture changes in effect direction. Since windows with larger effects can also deviate more from their fitted response in absolute terms, the dependence of seasonal deviation on effect magnitude was summarised as the proportion of variance in log(seasonal deviation) explained by log(magnitude) (R²) within each trait and omic layer. Back-solving and window analyses were performed for yield, brix and tartaric acid. These traits were chosen to span high (yield, brix) and intermediate (tartaric acid) clonal repeatability and to include predominantly scale-type (yield, brix) and crossover (tartaric acid) GEI.

To test whether the largest genomic and epigenomic effects occur in the same regions, we counted the windows ranked in the top 5% of effect magnitude in both layers (1,928 windows per layer). If the two layers were independent, about 96.4 shared windows would be expected by chance (1,928 × 1,928 / 38,558). Since windows with more features could have larger effects in both layers, overlap could arise from feature density alone. Significance was therefore assessed by permutation: in each of 1,000 permutations, the top-5% windows of each layer were reassigned at random among windows with similar numbers of features (20 quantile groups per layer), and the overlap was recounted. The one-sided p-value was the proportion of permutations with an overlap at least as large as that observed.

To check that window rankings were not driven by a single season, window statistics were recalculated nine times, each time leaving out one year, and the resulting rankings were compared with the full nine-year rankings using Spearman correlation and the retention of the 100 highest-ranked windows.

## Results

### Genomic and epigenomic relationships among clones are weakly structured

After filtering, the genomic dataset comprised 972,674 SNPs and the epigenomic dataset comprised 455,847 methylation features. PCA of the SNP-derived GRM showed relatively limited differentiation, with the first two principal components explaining 5.0% of the total variance (Figure 1a). A small number of clones were separated from the main cluster, primarily along PC2. The SNP minor allele frequency distribution was bimodal (Supplementary Figure 1a), with an enrichment of rare alleles alongside a second concentration of variants at intermediate frequencies. Rare SNPs (MAF ≤ 0.1) comprised 10.0% (97,465) of all markers and were broadly distributed across clones, with each variant observed in a median of 13% of individuals. Across the filtered SNP set, a mean of 56% of genotype calls per clone were heterozygous.

**Figure 1.**
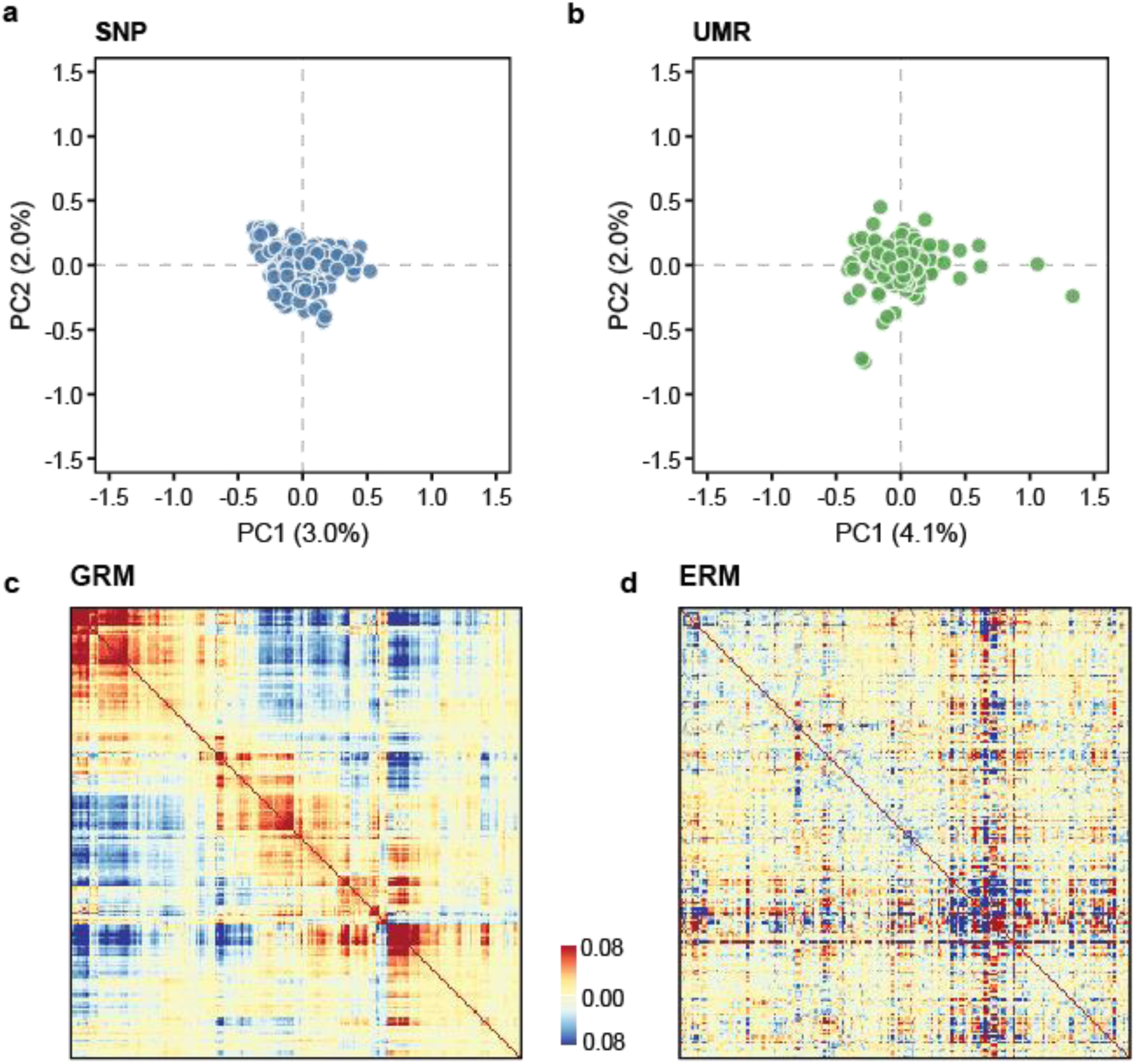
Molecular characterisation of genomic and epigenomic relationships in Riesling clonal population. **(a)** Principal component (PC) analysis of the SNP-derived genomic relationship matrix (GRM), showing relatively limited differentiation among clones, with PC1 and PC2 explaining 3.0% and 2.0% variance, respectively. **(b)** PC analysis of the orthogonalised epigenomic relationship matrix (ERM) used in all models, with PC1 and PC2 explaining 4.1% and 2.0% of the variance, respectively. **(c)** Heatmap of the GRM and **(d)** heatmap of the orthogonalized ERM, with red indicating higher relatedness and blue indicating lower relatedness.

Methylation Shannon entropy varied widely across loci (mean = 2.93, range = 0.04–4.20; maximum possible value log^2^ 20 ≈ 4.32), indicating considerable variation in methylation state among clones. The entropy distribution was bimodal within the higher range, with many loci showing moderate to high variability in methylation state. The GRM and non-orthogonalized ERM were strongly collinear (off-diagonal r = 0.71; Supplementary Figure S2a). Non- orthogonalized methylation features showed much stronger structure, with PC1 and PC2 together accounting for 55% of the variance (Supplementary Figure S2b). When the non- orthogonalised ERM was fitted alongside the GRM, its variance component was consistently estimated at the boundary, indicating redundancy due to collinearity between the matrices. To reduce this redundancy, the ERM was orthogonalized, and following this, the GRM (Figure 1c) and orthogonalised ERM (Figure 1d) were weakly correlated (off-diagonal r = -0.22). PCA of the orthogonalized ERM showed limited differentiation, similar to the GRM, with PC1 and PC2 explaining 4.1% and 2.0% of the variance (Figure 1b). Neither genomic nor epigenomic relationships were associated with clone passport data. Region of selection explained less than 4% of the variation in each relationship matrix (PERMANOVA p ≥ 0.47), generation and vineyard field each less than 1% (p ≥ 0.10, except vineyard field for the GRM, p = 0.049; Supplementary Table S3).

### Substantial clone level variation remains after environmental adjustment

After accounting for environmental effects, clone BLUEs showed clear variation across five yield and juice quality traits (Figure 2). Clone BLUEs ranged from 274.88 – 2084.41 g m⁻² for yield, 19.89 – 23.98 °Brix for brix, 5.47 – 7.63 g L⁻¹ for tartaric acid, 8.27 – 11.53 g L⁻¹ for total acidity, and 2.70 – 4.69 g L⁻¹ for malic acid. Clone-level BLUEs are substantially narrower than the corresponding raw phenotypic observations across years and fields (Figure 2). This is consistent with a considerable proportion of the observed phenotypic variation arising from environmental and other non-clonal sources. Despite this narrowing, clonal differences remained reproducible, where repeatability was highest for yield (H^2^ = 0.79) and brix (H^2^ = 0.70), intermediate for tartaric and malic acid (H^2^ = 0.45 for both), and lowest for total acidity (H^2^ = 0.33; Figure 2).

**Figure 2.**
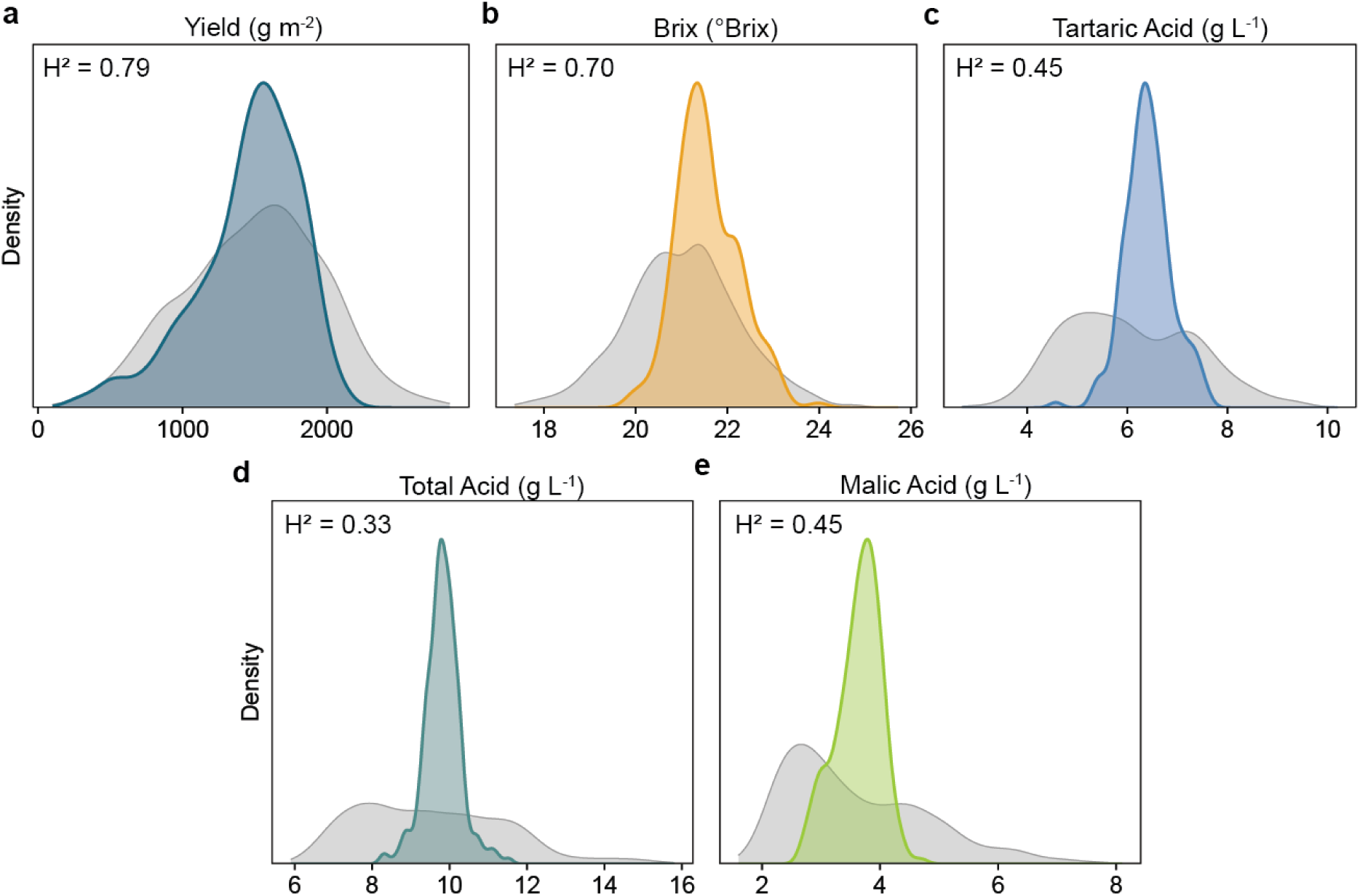
Phenotypic variation and clonal repeatability of yield and juice quality traits. Density distributions of raw observations across years and fields (grey) and clone best linear unbiased estimators (BLUEs; coloured) for (a) yield (g m^−2^), (b) brix (°Brix), (c) tartaric acid (g L^−1^), (d) total acidity (g L^−1^) and (e) malic acid (g L^−1^). Clonal repeatability (*H*^2^) is shown in the upper-left corner of each panel.

### Clonal variance is largely genomic when clone-by-year interaction is not modelled

In models without an explicit GEI term, clonal repeatability in the independent-clone model ranged from H^2^ = 0.33 for total acidity to H^2^ = 0.79 for yield. The kernel-based models recovered 93–107% of the clonal variance captured by this reference model and changed clonal repeatability by no more than H^2^ = 0.011 for any trait (Figure 3a; Supplementary Table S4). Partitioning the clonal variance captured by the multi-kernel (GRM + ERM) model showed that the genomic component accounted for the majority of clonal variance for four of the five traits, whereas Brix showed a slightly larger epigenomic contribution of 53% (Figure 3b). Relative to total phenotypic variance, the variance of the clonal main effect in these models, which assume no clone-by-year interaction, was generally modest and varied substantially among traits, from 1–2% for total acidity and malic acid to 32% for yield (Figure 3c). Environmental variance (year and year × field) was the largest non-residual component for all traits, ranging from 34% for yield to 85% for total acidity, and exceeded the residual variance for the three acid traits. Residual variance ranged from 14% for total acidity and malic acid to 43% for Brix.

**Figure 3.**
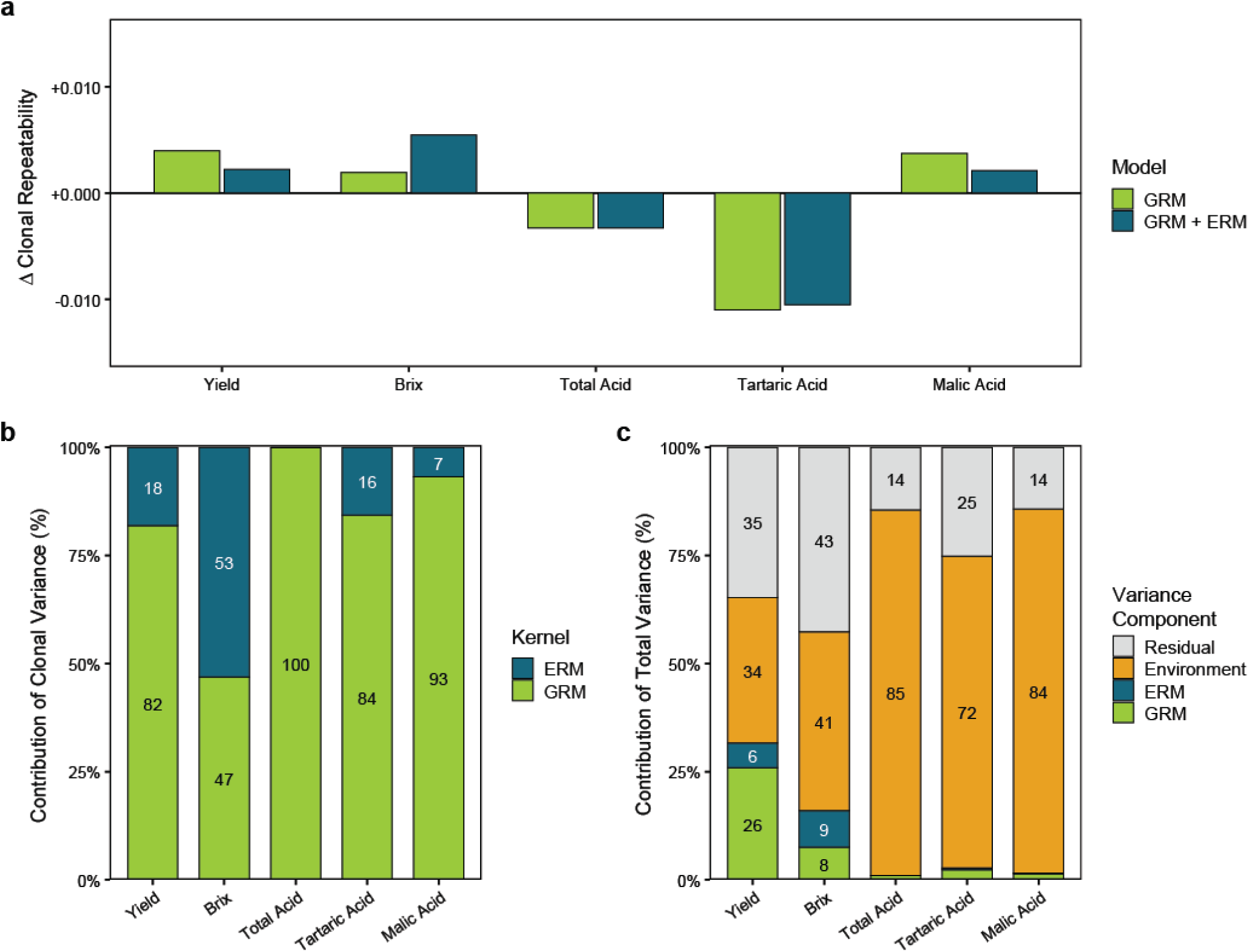
Clonal repeatability and total variance across traits. **(a)** Change in clonal repeatability relative to a reference model in which clones were treated as independent (absolute values of clonal repeatability are shown in Figure 2). The zero line indicates no change relative to the reference model. Bars show the genomic model (GRM; green) and the multi-kernel model (GRM + ERM; blue). (b) Partitioning of clonal variance between the genomic (GRM) and epigenomic (ERM) kernels in the multi-kernel model, expressed as a percentage of total clonal variance. (c) Partitioning of total phenotypic variance into genomic (GRM), epigenomic (ERM), environmental, and residual components in the multi-kernel model. Environmental variance comprises year and year × field effects. Traits are yield (g m⁻²), Brix (°Brix), total acidity (g L⁻¹), tartaric acid (g L⁻¹), and malic acid (g L⁻¹).

### Genomic and epigenomic relationship matrices capture different shares of clone-by-year variance

Clone-by-year effects captured by the genomic and epigenomic relationship matrices accounted for a substantial proportion of the phenotypic variance remaining after fixed year and field effects (hereafter, total variance) for all five traits (Figure 4). Combined Year:GRM and Year:ERM components explained 33.3–50.5% of total variance, with the largest contributions observed for Brix (50.5%) and berry yield (49.0%), and the smallest for total acidity (33.3%). The relative contributions of the two omic layers were trait dependent. The variance associated with Year:ERM exceeded that of Year:GRM for four of the five traits, with the strongest difference observed for yield (31.3% vs. 17.7%). More comparable contributions were observed for brix (27.5% vs. 23.0%), tartaric acid (18.3% vs. 16.5%), and malic acid (25.6% vs. 21.6%). In contrast, for total acidity, Year:GRM contributed approximately twice the variance of Year:ERM (22.3% vs. 11.0%).

**Figure 4.**
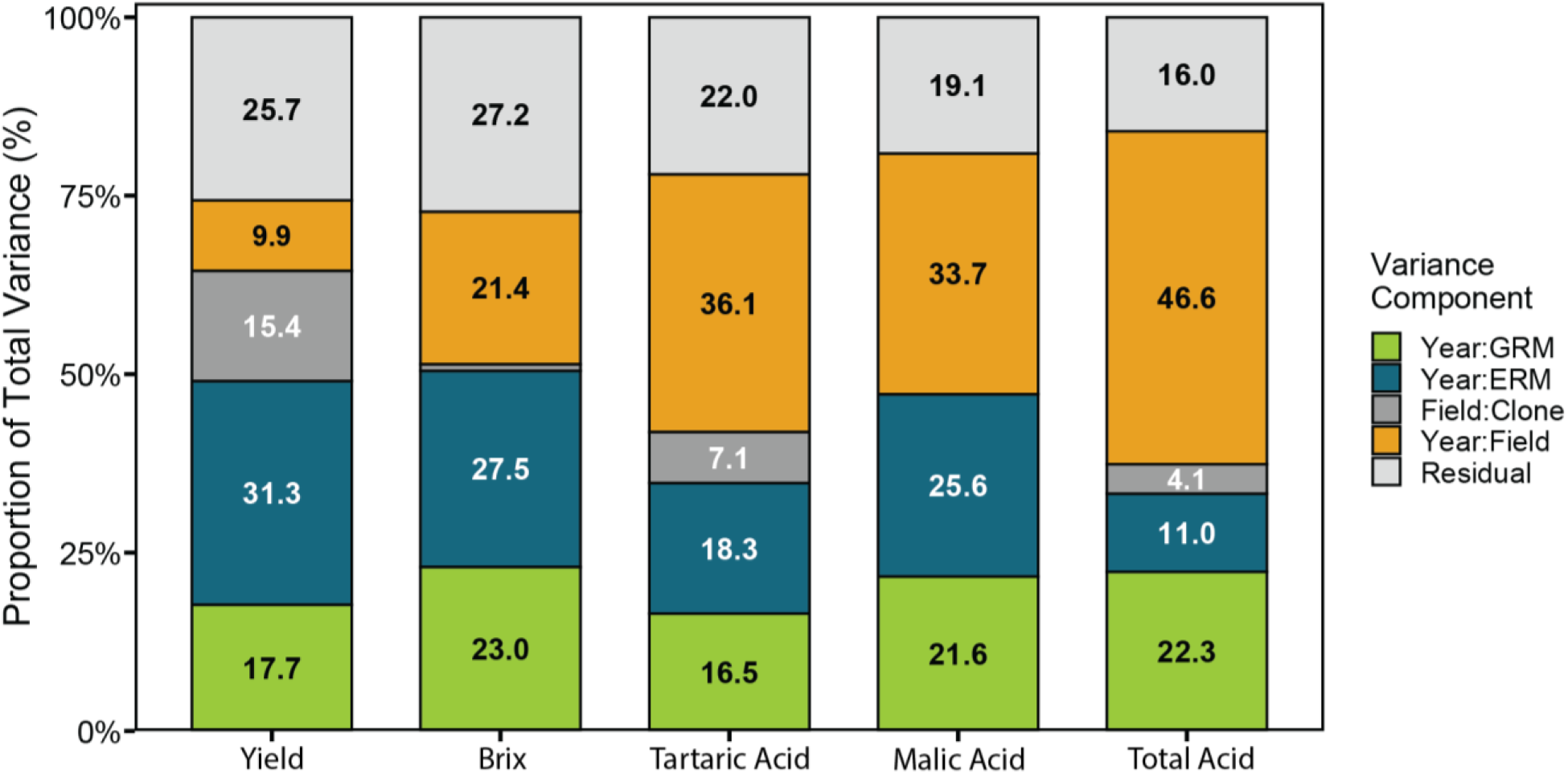
Partitioning of clone-by-environment variance across genomic and epigenomic layers. Variance decomposition from multi-kernel mixed models including GEI terms for yield (g m⁻²), brix (°Brix), tartaric acid (g L⁻¹), malic acid (g L⁻¹), and total acidity (g L⁻¹). Variance components are expressed as a percentage of the summed variance of all random effects, which excludes the fixed year and field effects (values are therefore not directly comparable with Figure 3c), and include genomic clone-by-year interaction (Year:GRM), epigenomic clone-by- year interaction (Year:ERM), clone-by-field interaction (Field:Clone), environmental year-by- field interaction (Year:Field), and residual variance (Residual). Values within bars indicate the percentage of this total attributed to each component.

The year-by-field component also varied substantially among traits, increasing from 9.9% for yield to 46.6% for total acidity. This ordering mirrored clonal repeatability in reverse, whereby the year-by-field share was smallest for the traits with the highest repeatability (yield, *H*^2^ = 0.79; Brix, *H*^2^ = 0.70; Figure 2), intermediate for malic and tartaric acid (33.7% and 36.1%; both *H*^2^ = 0.45) and largest for total acidity (*H*^2^ = 0.33). Clone-by-field variance was substantially smaller than the clone-by-year components, although its contribution was more pronounced for yield (15.4%) than for the other traits. Residual variance accounted for 16.0– 27.2% of total variance and was highest for brix and yield and lowest for total acidity.

### Genomic and epigenomic layers capture distinct patterns of clone re-ranking across years

The structure of clone-by-year effects differed between the genomic and epigenomic layers, with the extent and pattern of re-ranking varying among traits (Figure 5). For yield and brix, epigenomic correlations were consistently positive across years, indicating that rank of clone performance was largely conserved across seasons and that GEI was predominantly associated with changes in effect size. Genomic correlations for yield were also predominantly positive, although 2017 was negatively correlated with most other years, indicating pronounced re- ranking associated with this year. For brix, genomic correlations were weaker and more variable across years than their epigenomic counterparts. Tartaric acid and total acidity showed more complex GEI in both omic layers, with positive and negative between-year correlations indicating substantial re-ranking among years. For malic acid, genomic correlations were predominantly weak and positive, whereas epigenomic correlations showed greater variation among years.

**Figure 5.**
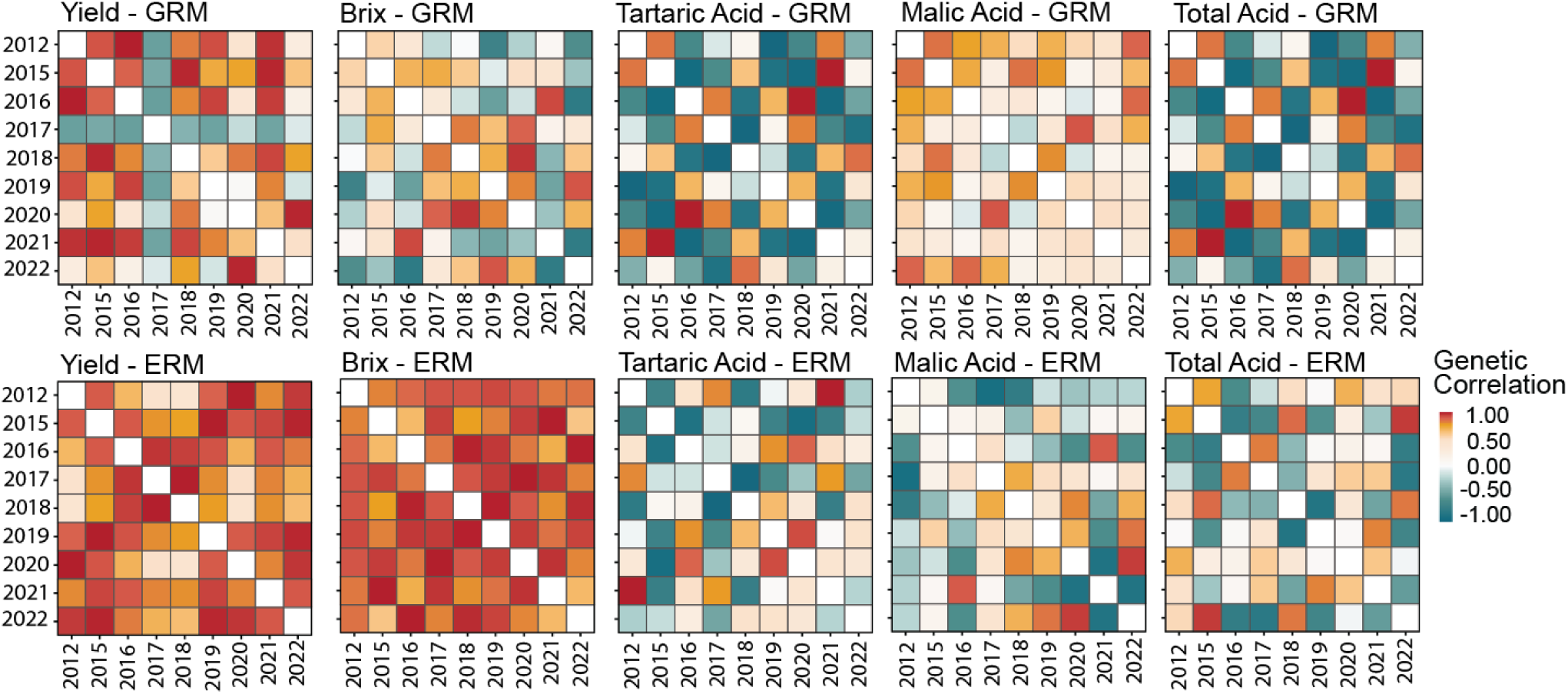
Between-year correlations of genomic and epigenomic clone effects. Correlations among years derived from factor-analytic covariance matrices fitted to genomic (GRM; upper row) and epigenomic (ERM; lower row) clone-by-year effects for yield, brix, tartaric acid, malic acid, and total acidity. Positive correlations (red) indicate greater maintenance of clone ranking between years, whereas negative correlations (blue) indicate greater re-ranking of clones between years. Diagonal cells are left blank.

### Large-effect genomic and epigenomic windows show limited positional overlap

Window-level effect magnitudes derived from back-solved locus effects were distributed broadly across the genome for both omic layers, with a small number of windows showing markedly larger trait effects (Figure 6). However, the highest-effect windows showed limited positional overlap between the genomic and epigenomic layers. Of the 1,928 windows comprising the top 5% of effect magnitude in each layer, 85 were shared for yield, 86 for Brix, and 92 for tartaric acid, compared with 96.4 expected under positional independence. For all three traits, the observed overlap did not exceed that expected under a permutation null that preserved differences in feature density among windows (p = 0.50, 0.59, and 0.17 for yield, brix, and tartaric acid, respectively).

**Figure 6.**
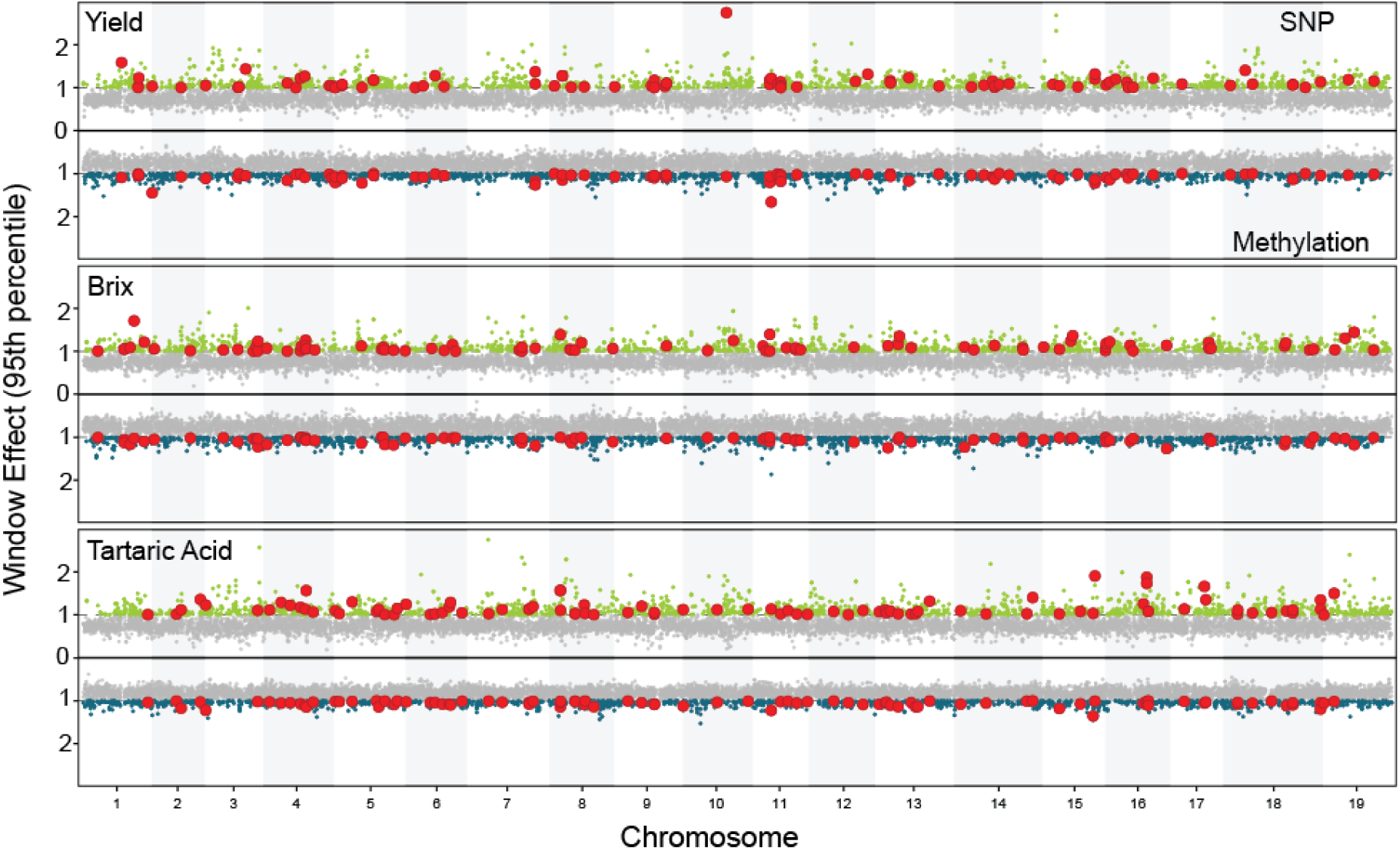
Genome-wide distribution of back-solved genomic and epigenomic window effects. Mirror plots of window-level effect magnitudes for SNPs (green; upper half) and methylation features (teal; lower half, inverted axis) across chromosomes for yield, brix, and tartaric acid. Back-solved locus effects were aggregated into 25 kb sliding windows with a 10 kb step and summarised as the mean absolute effect per feature, averaged across years. Values within each omic layer were scaled by the corresponding 95th percentile, such that a value of 1 represents the top 5% threshold. Grey points indicate windows below this threshold, coloured points indicate windows within the top 5% of their respective layer, and red points identify windows within the top 5% of both layers. Values greater than 3 are truncated for visualisation.

Window rankings were robust to leave-one-year-out analysis, with strong agreement between the full-data and reduced rankings (Spearman ρ ≥ 0.93) and 64–100% retention of the top 100 windows. The lowest retention occurred when 2020 was omitted from the epigenomic analysis of tartaric acid (Supplementary Table S5).

### Effect magnitude is more strongly associated with seasonal deviation in the genomic layer

The relationship between window effect magnitude and seasonal deviation, the departure of a window’s yearly effect from its fitted seasonal response, differed between the genomic and epigenomic layers (Figure 7). In the genomic layer, larger-effect windows tended to deviate more from their fitted seasonal response, with modest positive correlations for yield, brix, and tartaric acid (Spearman ρ = 0.35, 0.31, and 0.26, respectively). In the epigenomic layer, this association was negligible for yield (ρ = 0.07) and brix (ρ = −0.01) and weak for tartaric acid (ρ = 0.21). Across all traits and layers, effect magnitude accounted for no more than 13% of the variation in seasonal deviation.

**Figure 7.**
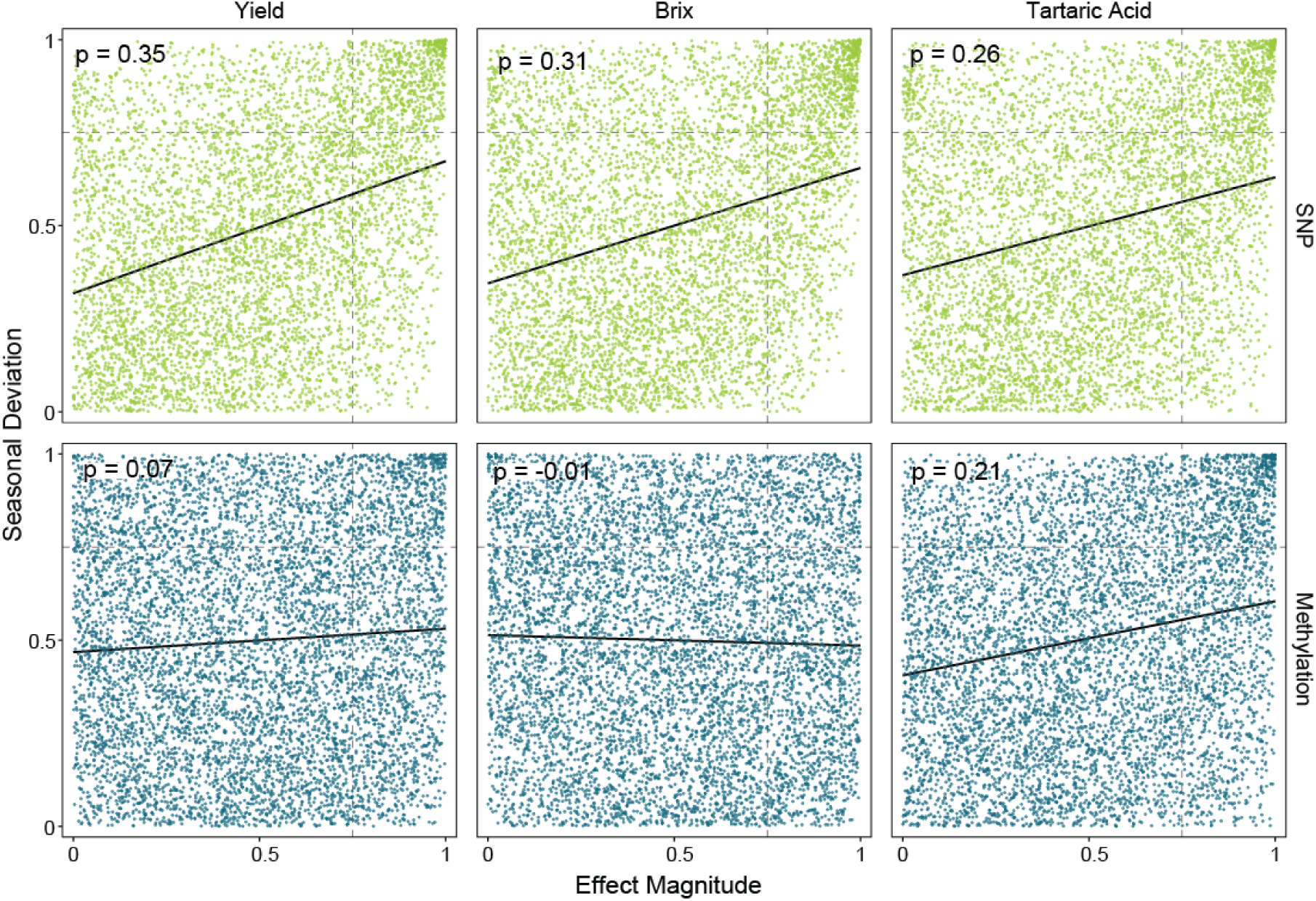
Relationship between window effect magnitude and seasonal deviation. Each point represents a 25 kb window for yield, brix, or tartaric acid in the genomic (SNP; upper row) and epigenomic (Methylation; lower row) layers. Effect magnitude and seasonal deviation are rank-transformed within each trait and omic layer. Seasonal deviation was quantified as the root mean square deviation of yearly window effects from their fitted regression on the yearly environmental index. Dashed lines indicate the upper quartile of each axis, black lines show linear fits, and values indicate Spearman’s ρ. For display purposes a random 25% of windows are shown; correlations were computed from all windows.

The genomic layer also showed an enrichment of windows combining large effects with high seasonal deviation. Windows in the upper quartile for both effect magnitude and seasonal deviation occurred 1.8- to 2.0-fold more frequently than expected under independence, whereas windows combining large effects with low seasonal deviation were under-represented (0.69- to 0.74-fold). This pattern was weaker or absent in the epigenomic layer, where enrichment of high-magnitude, high-deviation windows ranged from 1.1- to 1.5-fold.

## Discussion

### Clonal relationships show limited structure and no provenance signal

Genomic relationships among the Riesling clones showed limited structure, with the first two principal components explaining only 5% of the variance, consistent with their shared varietal background. Although the unadjusted methylation features showed stronger structure, the ERM was also weakly structured after orthogonalization with respect to genomic structure. Together, these results suggest that the substantial epigenomic contribution to clone-by-year variance reported here is unlikely to be driven primarily by broad genetic or epigenomic groupings among clones.

Neither genomic nor epigenomic relationships reflected recorded provenance. This contrasts with Pinot noir clones, for which relationships inferred from methylation polymorphisms closely matched somatic SNP-based relatedness and recorded provenance (Callipo et al., 2026). The absence of a provenance signal in either relationship matrix suggests that the course, region-level records of this historic Riesling collection may not adequately capture clonal ancestry. Methodological differences may also contribute to the contrasting findings, whereby Callipo et al. (2026) used whole-genome long-read sequencing and found concordance mainly in the CG and CHG contexts, whereas UMR-seq profiles unmethylated regions without resolving sequence context (Putland et al., 2026).

### Clone-by-year variance is primarily associated with epigenomic variation

In models without explicit GEI terms, adding the ERM had little effect on clonal repeatability, and the genomic kernel accounted for most of the partitioned clonal variance for four of the five traits (Figure 3). This is likely because each clone was phenotyped across multiple seasons, so clone means were already estimated with reasonable precision, and relationships among clones were relatively weak in both matrices, providing little additional information from related clones. Molecular relationship matrices may be more informative for clones with fewer records, such as newly selected material.

In contrast, in the GEI models, the epigenomic kernel accounted for a substantial proportion of clone-by-year variance, exceeding the genomic contribution for four of the five traits (Figure 4). This suggests that epigenomic relationships captured differences in how clones responded to individual seasons that were less apparent in their average performance across years. Previous studies have reported that integrating additional omic layers can increase variance capture relative to genomic data, although the magnitude and consistency of these gains are context dependent (Robinson et al., 2025b; Montesinos-López et al., 2025). In perennial ryegrass, integrating DNA methylation with genomic and transcriptomic data explained more phenotypic variance than any single-omic model (Malinowska et al., 2022). The present results extend this by showing that, in a clonal population, the contribution of methylation may be expressed primarily through GEI rather than through clonal main effects.

### Genomic and epigenomic layers capture distinct patterns of clone-by-year response

The relative contributions of genomic and epigenomic relationships to clone-by-year variance differed among traits. The epigenomic contribution exceeded the genomic contribution for four of the five traits, with the largest difference observed for yield (31.3% versus 17.7% of total variance). Total acidity showed the opposite pattern, with approximately twice as much variance attributed to the genomic component (22.3% versus 11.0%; Figure 4). Both kernels accounted for clone-by-year variance across all traits, suggesting that each captured information about differences in clonal responses among years. However, this statistical partitioning does not establish that the two layers represent separate biological mechanisms. Beyond these clone-specific responses, the year-by-field component accounted for 9.9% of total variance in yield to 46.6% in total acidity, indicating substantial variation among field– year combinations.

The distinction between the two omic layers was also evident in their between-year correlation structures (Figure 5). For yield and brix, epigenomic correlations were consistently positive across years, indicating relatively conserved clone rank performance. Genomic correlations for yield were also largely positive, although 2017 showed negative correlations with most other years, indicating substantially greater re-ranking associated with this year. For tartaric acid and total acidity, both layers showed a broader range of positive and negative correlations, consistent with greater re-ranking across years, although the specific patterns differed between layers. These results show that similar amounts of GEI variance can mask different patterns of response across environments and that genomic and epigenomic relationships do not necessarily describe the same clone-by-year structure. Individual between-year correlations should nevertheless be interpreted cautiously because several year-specific variances in the factor-analytic models were estimated close to zero, such that some correlations are more strongly determined by the fitted covariance structure.

Previous studies have demonstrated the value of multi-omic and multi-environment genomic prediction models for capturing GEI in perennials (Malinowska et al., 2022; Jung et al., 2022, 2025; Brault et al., 2024), but direct comparisons of omic-specific GEI covariance structures remain limited. By modelling genomic and epigenomic clone-by-year effects separately, the present analysis shows that the two layers differ not only in their contribution to GEI variance but also in the patterns of clone response they capture across years. Whether these differences improve prediction in unobserved environments remains an important question for future work.

### Genomic and epigenomic window effects differ in their distribution and temporal behaviour

The windowed analysis suggested that the strongest genomic and epigenomic effects were distributed differently across the genome. For yield, brix, and tartaric acid, overlap between the top 5% of windows in each layer was no greater than expected by chance (Figure 6), indicating that the two layers did not preferentially identify the same regions. This does not establish biological independence between genomic and epigenomic variation. Orthogonalization of the ERM accounted for genome-wide genomic structure but does not necessarily remove local associations between sequence and methylation variation. Consequently, methylation-derived window effects may still reflect underlying sequence variation, even where the strongest effects do not coincide between layers.

Window effect magnitude provided limited information about seasonal deviation, with the relationship differing between omic layers (Figure 7). In the genomic layer, larger-effect windows tended to deviate more from their fitted seasonal response, and windows with both high magnitude and high deviation were enriched relative to expectation. This association was weaker or absent in the epigenomic layer. Effect magnitude and seasonal deviation therefore provide complementary information, distinguishing the size of a window’s effect from how closely it follows the fitted seasonal pattern. For breeding, considering both characteristics could help distinguish large effects that track the overall seasonal response from those with more season-specific behaviour. Seasonal deviation was calculated from absolute effects and therefore does not distinguish changes in effect magnitude from reversals in direction. Between-year correlations provide a complementary view of crossover patterns (Figure 5). Functional validation will be required to establish whether the identified windows contain causal sequence or regulatory variants underlying these response patterns.

### Suitability of UMR-seq for population-scale analyses in grapevine

Population-scale analyses of clonal GEI require molecular profiles for hundreds of clones, a scale that remains challenging for whole-genome methylation assays. In grapevine, base- resolution methylation profiling has so far been restricted to comparatively small panels, such as 23 Pinot noir clones sequenced using long reads (Callipo et al., 2026), while earlier reduced- representation approaches sampled only a small fraction of the genome (Schmidt et al., 2025). UMR-seq offers an intermediate approach. Digestion with the methylation-dependent restriction enzymes depletes methylated DNA and concentrates sequencing on unmethylated regions (Putland et al., 2026). In plants, these regions are largely stable across vegetative tissues and mark expressed genes and their cis-regulatory space (Crisp et al., 2020), directing sequencing towards regions likely to be functionally relevant. Libraries were prepared from 1 ng of digested DNA using half-volume reactions, with SNPs and methylation features recovered from the same reads. Together, these features enabled profiling of more than 200 clones in a single study, addressing two constraints on the use of epigenomic data in perennial crop breeding: cost per sample and the need for separate assays for different omic layers (Robinson et al., 2025b).

Several features of the present data are consistent with UMR-seq capturing structured variation at this scale. Before adjustment, the ERM was strongly correlated with the GRM (off-diagonal r = 0.71), consistent with findings from a population-scale UMR-seq analysis in wheat (Putland et al., 2026). Support for the genomic layer also comes from wheat, where SNP genotypes obtained from UMR-seq libraries were strongly correlated with those from an independent SNP array (Putland et al., 2026). This concordance indicates that UMR-targeted markers can recover broad genetic relationships despite sampling only part of the genome. The UMR-seq methylation profiles provide regional information for analysing clonal variation, complementing the sequence-context and haplotype-specific detail available from long-read methylation profiling (Callipo et al., 2026). Coverage-derived methylation features may also reflect underlying sequence variation and should therefore be interpreted as regional signals rather than direct measurements of individual cytosine methylation states. The present results demonstrate the utility of these profiles for characterising genomic and epigenomic contributions to clonal performance across years, supporting the wider application of UMR- seq to multi-omic predictive modelling.

## Limitations and future directions

The results presented here should be interpreted within the context of the study’s scope. First, the study represents a single Riesling clonal population, predominantly sourced from Germany, and phenotyped in a historical nursery with limited within-field and partial between-field replication. Repeated phenotyping across nine years and mixed models incorporating molecular relationships helped account for this unbalanced design, but validation in independent clonal populations will be required to establish the generality of the observed genomic and epigenomic patterns.

Second, the epigenomic profiles were measured at a single time point in 2021. Relating these profiles to phenotypes across multiple years therefore assumes that at least part of the methylation variation captured by the methylation data is sufficiently persistent to remain informative across seasons. Evidence for this is strongest for unmethylated regions, which remain stable across vegetative tissues in maize (Crisp et al., 2020). In Pinot noir clones, the CG and CHG methylation polymorphisms that distinguish clonal lineages showed frequency distributions consistent with mitotic maintenance (Callipo et al., 2026), but how the more dynamic component of the methylome behaves across time points and tissues and how this is captured by UMRseq is not known.

Finally, The window analyses provide candidates for further investigation, with their effects and seasonal behaviour remaining to be tested in independent populations and environments. In addition, the omic-by-year terms fitted in the mixed model partition GEI statistically but do not identify the environmental factors driving these interactions. Integrating measured environmental covariates into the multi-omic framework is therefore an important next step, enabling genomic and epigenomic response patterns to be linked directly to climatic or management variables and tested for their ability to predict performance in unobserved environments.

## Conclusions

This study integrated genome-wide SNP and DNA methylation data to partition clonal and clone-by-year variation in a Riesling grapevine population. Paired profiles from a single UMR- seq library per clone showed that genomic and epigenomic relationships among clones were weakly structured and unrelated to recorded provenance. In models without explicit GEI, incorporating epigenomic relationships had little effect on clonal repeatability. In contrast, when clone-by-year effects were modelled explicitly, the epigenomic layer captured a substantial proportion of GEI variance and exceeded the genomic contribution for the majority of traits. The two omic layers also captured distinct patterns of clone response across years, showing that similar contributions to GEI variance can represent different patterns of clone re- ranking. At the genomic level, large-effect genomic and epigenomic windows showed no greater positional overlap than expected by chance, while effect magnitude was only weakly associated with deviation from the seasonal response, and this relationship was stronger in the genomic layer. Together, these findings show that genomic and epigenomic relationships capture complementary components of year-to-year variation in clone performance, with the epigenomic contribution expressed primarily through clone-by-year variation. Extending this framework to independent clonal populations and incorporating measured environmental covariates will be important for determining whether these distinct omic signals can improve prediction of performance in unobserved environments.

## Author contributions

HR: conceptualisation, formal analysis, visualisation, writing – original draft. KPV-F: conceptualisation, supervision, interpretation, funding acquisition, writing – review and editing. MS: UMR-seq data processing pipeline, writing – review and editing. JF: DNA extraction. JH and UV: UMR-seq library preparation. PC: UMR-seq methodology, writing – review and editing. H-PP: statistical methodology, writing – review and editing. PC, CR-Z, SB and TS: interpretation, writing – review and editing.

## Acknowledgements

Vineyard and laboratory staff at Hochschule Geisenheim University

## References

1. Berger, S. L., Kouzarides, T., Shiekhattar, R., & Shilatifard, A. (2009). An operational definition of epigenetics. Genes & Development, 23, 781–783. 10.1101/gad.1787609

2. Bolger, A. M., Lohse, M., & Usadel, B. (2014). Trimmomatic: A flexible trimmer for Illumina sequence data. Bioinformatics, 30(15), 2114–2120. 10.1093/bioinformatics/btu170

3. Brault, C., Doligez, A., Le Cunff, L., Coupel-Ledru, A., Simonneau, T., Chiquet, J., This, P., & Flutre, T. (2021). Harnessing multivariate, penalized regression methods for genomic prediction and QTL detection of drought-related traits in grapevine. G3: Genes, Genomes, Genetics, 11(9), jkab248. 10.1093/g3journal/jkab248

4. Brault, C., Segura, V., This, P., Le Cunff, L., Flutre, T., François, P., Pons, T., Péros, J.-P., & Doligez, A. (2022a). Across-population genomic prediction in grapevine opens up promising prospects for breeding. Horticulture Research, 9, uhac041. 10.1093/hr/uhac041

5. Brault, C., Lazerges, J., Doligez, A., Thomas, M., Ecarnot, M., Roumet, P., & Segura, V. (2022b). Interest of phenomic prediction as an alternative to genomic prediction in grapevine. Plant Methods, 18(1), 108. 10.1186/s13007-022-00940-9

6. Brault, C., Segura, V., Roques, M., Lamblin, P., Bouckenooghe, V., Pouzalgues, N., Cunty, C., Breil, M., Frouin, M., & Garcin, L. (2024). Enhancing grapevine breeding efficiency through genomic prediction and selection index. G3: Genes, Genomes, Genetics, 14(4), jkae038. 10.1093/g3journal/jkae038

7. Broad Institute. (2019). Picard toolkit. GitHub repository. http://broadinstitute.github.io/picard/

8. Butler, D. G., Cullis, B. R., Gilmour, A. R., Gogel, B. J., & Thompson, R. (2023). ASReml-R reference manual version 4.2. VSN International Ltd.

9. Callipo, P., Schmidt, M., Strack, T., Robinson, H., Vasudevan, A., & Voss-Fels, K. P. (2025). Harnessing clonal diversity in grapevine: From genomic insights to modern breeding applications. Theoretical and Applied Genetics, 138, 196. 10.1007/s00122-025-04986-w

10. Callipo, P., Robinson, H., Schmidt, M., & Voss-Fels, K. P. (2026). A dual genome- methylome map of clonal evolution in grapevine. Genome Biology, 27, 221. 10.1186/s13059-026-04184-x

11. Carbonell-Bejerano, P., Royo, C., Torres-Pérez, R., Grimplet, J., Fernandez, L., Franco- Zorrilla, J. M., Lijavetzky, D., Baroja, E., Martínez, J., García-Escudero, E., Ibáñez, J., & Martínez-Zapater, J. M. (2017). Catastrophic unbalanced genome rearrangements cause somatic loss of berry color in grapevine. Plant Physiology, 175(2), 786–801. 10.1104/pp.17.00715

12. Cohen-Karni, D., Xu, D., Apone, L., Fomenkov, A., Sun, Z., Davis, P. J., Kinney, S. R. M., Yamada-Mabuchi, M., Xu, S.-Y., Davis, T., Pradhan, S., Roberts, R. J., & Zheng, Y. (2011). The MspJI family of modification-dependent restriction endonucleases for epigenetic studies. Proceedings of the National Academy of Sciences, 108(27), 11040–11045. 10.1073/pnas.1018448108

13. Cooper, M., & Messina, C. D. (2023). Breeding crops for drought-affected environments and improved climate resilience. The Plant Cell, 35(1), 162–186. 10.1093/plcell/koac321

14. Cooper, M., Voss-Fels, K. P., Messina, C. D., Tang, T., & Hammer, G. L. (2021). Tackling G × E × M interactions to close on-farm yield-gaps: Creating novel pathways for crop improvement by predicting contributions of genetics and management to crop productivity. Theoretical and Applied Genetics, 134(6), 1625–1644. 10.1007/s00122-021-03812-3

15. Crisp, P. A., Ganguly, D., Eichten, S. R., Borevitz, J. O., & Pogson, B. J. (2016). Reconsidering plant memory: Intersections between stress recovery, RNA turnover, and epigenetics. Science Advances, 2(2), e1501340. 10.1126/sciadv.1501340

16. Crisp, P. A., Marand, A. P., Noshay, J. M., Zhou, P., Lu, Z., Schmitz, R. J., & Springer, N. M. (2020). Stable unmethylated DNA demarcates expressed genes and their cis-regulatory space in plant genomes. Proceedings of the National Academy of Sciences, 117(38), 23991– 24000. 10.1073/pnas.2010250117

17. Putland, V., Robinson, H., Eglitis-Sexton, J. I. M., Yoshikawa, G. V., Hintzsche, J., Andrews, H., Andersen, S., Yoon, S., Tong, J., Mangila, L. M., Sree, A., Zanini, S., Schmidt, M., Zhang, Y., Snowdon, R. J., Golicz, A., Voss-Fels, K. P., Hickey, L. T., & Crisp, P. A. (2026). High-throughput sequencing of unmethylated DNA (UMR-seq) reveals extensive epigenetic variation in the hypomethylome of diverse plant species. bioRxiv. 10.64898/2026.10.29.762724

18. Cullis, B. R., Smith, A. B., & Coombes, N. E. (2006). On the design of early generation variety trials with correlated data. Journal of Agricultural, Biological, and Environmental Statistics, 11(4), 381–393. 10.1198/108571106X154443

19. Danecek, P., Auton, A., Abecasis, G., Albers, C. A., Banks, E., DePristo, M. A., Handsaker, R. E., Lunter, G., Marth, G. T., Sherry, S. T., McVean, G., & Durbin, R. (2011). The variant call format and VCFtools. Bioinformatics, 27(15), 2156–2158. 10.1093/bioinformatics/btr330

20. Danecek, P., Bonfield, J. K., Liddle, J., Marshall, J., Ohan, V., Pollard, M. O., Whitwham, A., Keane, T., McCarthy, S. A., Davies, R. M., & Li, H. (2021). Twelve years of SAMtools and BCFtools. GigaScience, 10(2), giab008. 10.1093/gigascience/giab008

21. Douhovnikoff, V., & Dodd, R. S. (2015). Epigenetics: A potential mechanism for clonal plant success. Plant Ecology, 216(2), 227–233. 10.1007/s11258-014-0430-z

22. Eberhart, S. A., & Russell, W. A. (1966). Stability parameters for comparing varieties. Crop Science, 6(1), 36–40. 10.2135/cropsci1966.0011183X000600010011x

23. Falconer, D. S., & Mackay, T. F. C. (1996). Introduction to quantitative genetics (4th ed.). Longman.

24. Feldmann, M. J., Piepho, H.-P., & Knapp, S. J. (2022). Average semivariance directly yields accurate estimates of the genomic variance in complex trait analyses. G3: Genes, Genomes, Genetics, 12(6), jkac080. 10.1093/g3journal/jkac080

25. Flutre, T., Le Cunff, L., Fodor, A., Launay, A., Romieu, C., Berger, G., Bertrand, Y., Terrier, N., Beccavin, I., Bouckenooghe, V., Roques, M., Pinasseau, L., Verbaere, A., Sommerer, N., Cheynier, V., Bacilieri, R., Boursiquot, J.-M., Lacombe, T., Laucou, V., … Doligez, A. (2022). A genome-wide association and prediction study in grapevine deciphers the genetic architecture of multiple traits and identifies genes under many new QTLs. G3: Genes, Genomes, Genetics, 12(7), jkac103. 10.1093/g3journal/jkac103

26. Illumina. (2019). Nextera XT DNA library prep reference guide (Document No. 15031942). Illumina, Inc.

27. Jung, M., Keller, B., Roth, M., Aranzana, M. J., Auwerkerken, A., Guerra, W., Al-Rifaï, M., Lewandowski, M., Sanin, N., Rymenants, M., Didelot, F., Dujak, C., Font i Forcada, C., Knauf, A., Laurens, F., Studer, B., Muranty, H., & Patocchi, A. (2022). Genetic architecture and genomic predictive ability of apple quantitative traits across environments. Horticulture Research, 9, uhac028. 10.1093/hr/uhac028

28. Jung, M., Quesada-Traver, C., Roth, M., Aranzana, M. J., Muranty, H., Rymenants, M., Guerra, W., Holzknecht, E., Pradas, N., Lozano, L., Didelot, F., Laurens, F., Yates, S., Studer, B., Broggini, G. A. L., & Patocchi, A. (2025). Integrative multi-environmental genomic prediction in apple. Horticulture Research, 12(2), uhae319. 10.1093/hr/uhae319

29. Lin, C. S., Binns, M. R., & Lefkovitch, L. P. (1986). Stability analysis: Where do we stand? Crop Science, 26(5), 894–900. 10.2135/cropsci1986.0011183X002600050012x

30. Lorenz, D. H., Eichhorn, K. W., Bleiholder, H., Klose, R., Meier, U., & Weber, E. (1995). Growth stages of the grapevine: Phenological growth stages of the grapevine (Vitis vinifera L. ssp. vinifera)—Codes and descriptions according to the extended BBCH scale. Australian Journal of Grape and Wine Research, 1(2), 100–103. 10.1111/j.1755-0238.1995.tb00085.x

31. Malinowska, M., Ruud, A. K., Jensen, J., Svane, S. F., Smith, A. G., Bellucci, A., Lenk, I., Nagy, I., Fois, M., Didion, T., Thorup-Kristensen, K., Jensen, C. S., & Asp, T. (2022). Relative importance of genotype, gene expression, and DNA methylation on complex traits in perennial ryegrass. The Plant Genome, 15(4), e20253. 10.1002/tpg2.20253

32. McKenna, A., Hanna, M., Banks, E., Sivachenko, A., Cibulskis, K., Kernytsky, A., Garimella, K., Altshuler, D., Gabriel, S., Daly, M., & DePristo, M. A. (2010). The Genome Analysis Toolkit: A MapReduce framework for analyzing next-generation DNA sequencing data. Genome Research, 20(9), 1297–1303. 10.1101/gr.107524.110

33. Montesinos-López, O. A., Montesinos-López, A., Mosqueda-González, B. A., Delgado- Enciso, I., Chavira-Flores, M., Crossa, J., Dreisigacker, S., Sun, J., & Ortiz, R. (2025). Genomic prediction powered by multi-omics data. Frontiers in Genetics, 16, 1636438. 10.3389/fgene.2025.1636438

34. Ocaña, J., Walter, B., & Schellenbaum, P. (2013). Stable MSAP markers for the distinction of Vitis vinifera cv Pinot Noir clones. Molecular Biotechnology, 55(3), 236–248. 10.1007/s12033-013-9675-3

35. Oksanen, J., Simpson, G. L., Blanchet, F. G., Kindt, R., Legendre, P., Minchin, P. R., O’Hara, R. B., Solymos, P., Stevens, M. H. H., Szoecs, E., & Wagner, H. (2026). vegan: Community ecology package (R package version 2.7-5). https://vegandevs.github.io/vegan/

36. Piepho, H.-P. (2023). An adjusted coefficient of determination (R²) for generalized linear mixed models in one go. Biometrical Journal, 65(7), 2200290. 10.1002/bimj.202200290

37. Poplin, R., Ruano-Rubio, V., DePristo, M. A., Fennell, T. J., Carneiro, M. O., Van der Auwera, G. A., Kling, D. E., Gauthier, L. D., Levy-Moonshine, A., Roazen, D., Shakir, K., Thibault, J., Chandran, S., Whelan, C., Lek, M., Gabriel, S., Daly, M. J., Neale, B., MacArthur, D. G., & Banks, E. (2018). Scaling accurate genetic variant discovery to tens of thousands of samples. bioRxiv, 201178. 10.1101/201178

38. Quinlan, A. R., & Hall, I. M. (2010). BEDTools: A flexible suite of utilities for comparing genomic features. Bioinformatics, 26(6), 841–842. 10.1093/bioinformatics/btq033

39. Robinson, H., Strack, T., Schmidt, M., Callipo, P., Nsibi, M., Schmid, J., Rühl, E., Piepho, H.-P., & Voss-Fels, K. P. (2025a). Exploring intra-varietal variation for complex traits in grapevine (Vitis vinifera L.). Theoretical and Applied Genetics, 138(12), 305. 10.1007/s00122-025-05088-3

40. Robinson, H., Robles-Zazueta, C. A., & Voss-Fels, K. P. (2025b). Accelerating perennial crop improvement via multi-omics-based predictive breeding. The Plant Genome, 18, e70058. 10.1002/tpg2.70058

41. Schmidt, M. (2026). UMR-seq processing pipeline (Version 1.0.0) [Computer software]. GitHub. https://github.com/HGU-Plant-Breeding/UMR-process-coverage; https://github.com/HGU-Plant-Breeding/UMR-process-SNP.

42. Schmidt, M., Strack, T., Andrews, H., Hickey, L. T., Crisp, P. A., & Voss-Fels, K. P. (2025). A new climate for genomic and epigenomic innovation in grapevine. Molecular Horticulture, 5, 44. 10.1186/s43897-025-00171-1

43. Seneviratne, S. I., Zhang, X., Adnan, M., Badi, W., Dereczynski, C., Di Luca, A., Ghosh, S., Iskandar, I., Kossin, J., Lewis, S., Otto, F., Pinto, I., Satoh, M., Vicente-Serrano, S. M., Wehner, M., & Zhou, B. (2021). Weather and climate extreme events in a changing climate. In V. Masson-Delmotte et al. (Eds.), Climate change 2021: The physical science basis. Contribution of Working Group I to the Sixth Assessment Report of the Intergovernmental Panel on Climate Change (pp. 1513–1766). Cambridge University Press. 10.1017/9781009157896.013

44. Smith, A., Cullis, B., & Thompson, R. (2001). Analyzing variety by environment data using multiplicative mixed models and adjustments for spatial field trend. Biometrics, 57(4), 1138– 1147. 10.1111/j.0006-341X.2001.01138.x

45. van Leeuwen, C., Sgubin, G., Bois, B., Ollat, N., Swingedouw, D., Zito, S., & Gambetta, G. A. (2024). Climate change impacts and adaptations of wine production. Nature Reviews Earth & Environment, 5(4), 258–275. 10.1038/s43017-024-00521-5

46. VanRaden, P. M. (2008). Efficient methods to compute genomic predictions. Journal of Dairy Science, 91(11), 4414–4423. 10.3168/jds.2007-0980

47. Varela, A., Ibañez, V. N., Alonso, R., Zavallo, D., Asurmendi, S., Gomez Talquenca, S., Marfil, C. F., & Berli, F. J. (2021). Vineyard environments influence Malbec grapevine phenotypic traits and DNA methylation patterns in a clone-dependent way. Plant Cell Reports, 40(1), 111–125. 10.1007/s00299-020-02617-w

48. Varela, A. M., Marfil, C. F., Gómez Talquenca, S., Fontana, A. R., Asurmendi, S., Buscema, F., & Berli, F. J. (2024). Three-year study of DNA cytosine methylation dynamics in transplanted Malbec grapevines. Plant Science, 342.

49. Vasimuddin, M., Misra, S., Li, H., & Aluru, S. (2019). Efficient architecture-aware acceleration of BWA-MEM for multicore systems. In 2019 IEEE International Parallel and Distributed Processing Symposium (IPDPS) (pp. 314–324). IEEE. 10.1109/IPDPS.2019.00041

50. Velt, A., Frommer, B., Blanc, S., Holtgräwe, D., Duchêne, É., Dumas, V., Grimplet, J., Hugueney, P., Kim, C., Lahaye, M., Matus, J. T., Navarro-Payá, D., Orduña, L., Tello-Ruiz, M. K., Vitulo, N., Ware, D., & Rustenholz, C. (2023). An improved reference of the grapevine genome reasserts the origin of the PN40024 highly homozygous genotype. G3: Genes, Genomes, Genetics, 13(5), jkad067. 10.1093/g3journal/jkad067

51. Vitezica, Z. G., Varona, L., & Legarra, A. (2013). On the additive and dominant variance and covariance of individuals within the genomic selection scope. Genetics, 195(4), 1223–1230. 10.1534/genetics.113.155176

52. Vondras, A. M., Minio, A., Blanco-Ulate, B., Figueroa-Balderas, R., Penn, M. A., Zhou, Y., Seymour, D., Ye, Z., Liang, D., Espinoza, L. K., Anderson, M. M., Walker, M. A., Gaut, B., & Cantu, D. (2019). The genomic diversification of grapevine clones. BMC Genomics, 20(1), 972. 10.1186/s12864-019-6211-2

53. Xiao, H., Wang, Y., Liu, W., Shi, X., Huang, S., Cao, S., Long, Q., Wang, X., Liu, Z., Xu, X., Peng, Y., Wang, P., Jiang, Z., Riaz, S., Walker, A. M., Gaut, B. S., Huang, S., & Zhou, Y. (2025). Impacts of reproductive systems on grapevine genome and breeding. Nature Communications, 16, 2031. 10.1038/s41467-025-56817-7

54. Xie, H., Konate, M., Sai, N., Tesfamicael, K. G., Cavagnaro, T., Gilliham, M., Breen, J., Metcalfe, A., Stephen, J. R., De Bei, R., Collins, C., & Lopez, C. M. R. (2017). Global DNA methylation patterns can play a role in defining terroir in grapevine (Vitis vinifera cv. Shiraz). Frontiers in Plant Science, 8, 1860. 10.3389/fpls.2017.01860

55. Yadav, S., Wei, X., Joyce, P., Atkin, F., Deomano, E., Sun, Y., Nguyen, L. T., Ross, E. M., Cavallaro, T., Aitken, K. S., Hayes, B. J., & Voss-Fels, K. P. (2021). Improved genomic prediction of clonal performance in sugarcane by exploiting non-additive genetic effects. Theoretical and Applied Genetics, 134(7), 2235–2252. 10.1007/s00122-021-03822-1

56. Zhang, H., Lang, Z., & Zhu, J. K. (2018). Dynamics and function of DNA methylation in plants. Nature Reviews Molecular Cell Biology, 19(8), 489–506. 10.1038/s41580-018-0016-z

