## Supplementary Methods for "Genomic and Epigenomic Relationships Capture Different Patterns of Genotype-by-Environment Interaction in Grapevine Clones"

**Non-additive genomic and epigenomic relationship matrices**

A genomic dominance relationship matrix (GRM-D) was constructed following the Natural and Orthogonal InterActions (NOIA) framework (Vitezica et al., 2013). Dominance deviations were coded as $-{2p}_{j}^{2}$, $2p_{j}q_{j}$, $-{2p}_{j}^{2}$ for genotypes 0, 1, and 2 respectively, where $p_{j}$ and $q_{j}$ are the major and minor allele frequencies at locus $j$. The resulting deviation matrix $D$ was mean-centred and the dominance relationship matrix computed as $D=\frac{{DD}^{T}}{p}$, trace-normalised such that the mean diagonal element equals one. Additive-by-additive (GRM-I), additive-by-dominance (GRM-AD), and dominance-by-dominance (GRM-DD) epistatic interaction matrices were constructed as the trace-normalised Hadamard products $G$∘$G$, $G$∘$D$, and $D$∘$D$ respectively. A non-additive epigenomic relationship matrix (ERM-N) was constructed from the orthogonalized methylation features. Each feature was squared to capture second-order effects, the squared values were mean-centred, and the matrix computed as ERM-N $={{MM}^{T}}/q$ where $M$ is the centred matrix of squared features, trace-normalised as above. Since coverage-based methylation data do not resolve allelic state, this term captures curvature in the methylation–phenotype relationship.
