## Supplementary Figures for "Genomic and Epigenomic Relationships Capture Different Patterns of Genotype-by-Environment Interaction in Grapevine Clones"

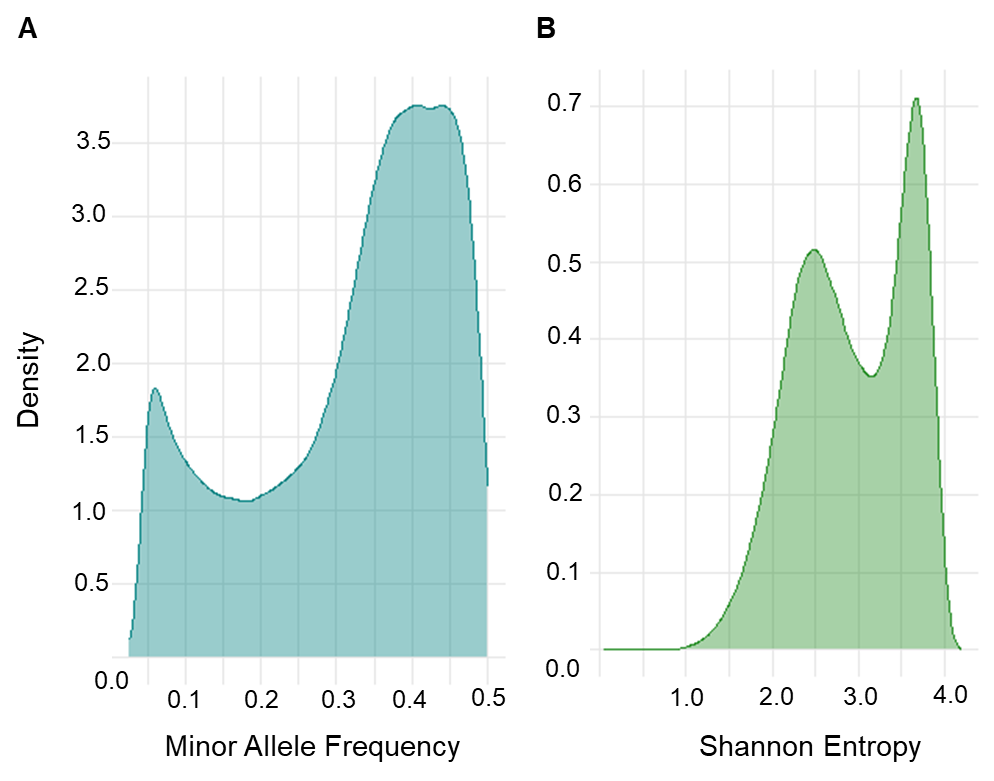

**Supplementary Figure 1. Distribution of SNP minor allele frequency and methylation Shannon entropy of Riesling clones. (a)** Density distribution of SNP minor allele frequency, showing a bimodal pattern. **(b)** Density distribution of methylation Shannon entropy across loci, showing a bimodal pattern within the higher entropy range.

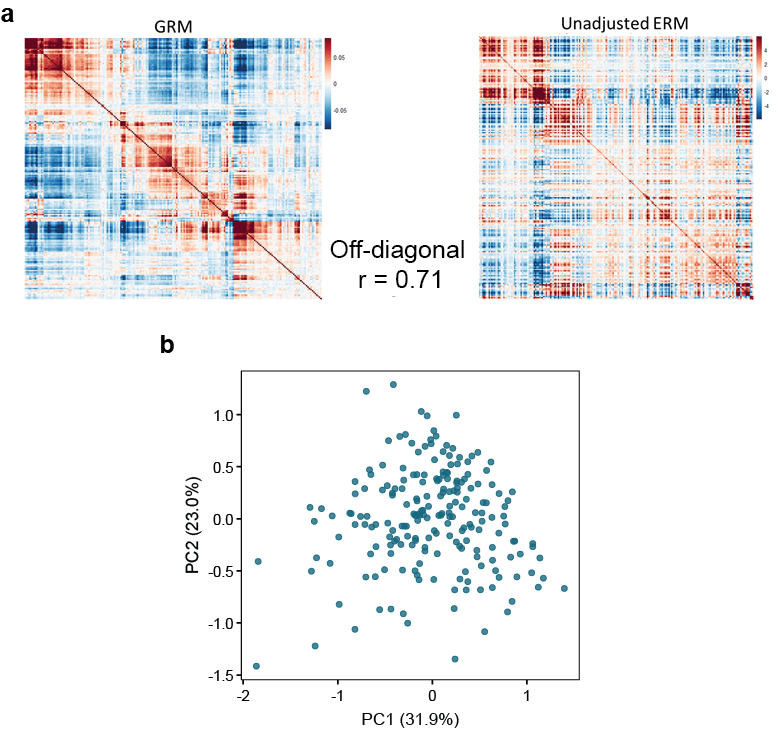

**Supplementary Figure 2. Collinearity between genomic and epigenomic relationship matrices.**
**(a)** Heatmaps of the genomic relationship matrix (GRM; left) and the unadjusted epigenomic relationship matrix (ERM; right), showing strong concordance in overall structure, with high correlation of off-diagonal elements (r = 0.71). **(b)** Principal component analysis of unadjusted ERM, showing differentiation among clones, with PC1 and PC2 explaining 31.9% and 23.0% variance, respectively.
